# Determining structures of biomacromolecular complexes from ambiguous NMR restraints and cryoEM data

**DOI:** 10.1101/2020.03.27.990309

**Authors:** Reto Walser, Alexander G. Milbradt

## Abstract

Integrated structural biology aims at combining different techniques to tackle challenging systems. Where individual techniques are not delivering structures of suitable quality, harnessing the strengths of various methods can often overcome this problem. X-ray crystallography and NMR have been the two most widely applied structural biology disciplines. In recent years cryoelectron microscopy (cryoEM) has become ever more powerful and is now capable of providing structures at resolutions comparable to those common in X-ray crystallography. Unfortunately, both NMR and cryoEM have inherent limitations on the system under study. However, the two techniques can be considered somewhat complementary as NMR has an upper and cryoEM a lower molecular weight (MW) limit. Here, we present a joint NMR and cryoEM methodology for the determination of biomacromolecular structures at the boundary region between the MW limits of the two techniques. The method relies on measuring chemical shift perturbations, which is the most readily accessible NMR parameter for characterizing the interaction of biomacromolecular complexes. Low-resolution cryoEM information yields global information on the shape of the complex and is used for complementing the local NMR data. We have successfully applied this method to the model system histidine-containing phosphoprotein (HPr) in complex with the glucose-specific acceptor protein IIA^Glc^ from *Escherichia coli*.

---

Modern biology calls for understanding biological processes at atomic detail. X-ray crystallography and nuclear magnetic resonance (NMR) spectroscopy have been successfully used to this end for many decades. In the past years also cryoelectron microscopy (cryoEM) has advanced to become a *bona fide* technique for delivering high-resolution structures.

The combination of the above methods, often complemented by other biophysical methods (*e.g.* mass spectrometry (MS), solution scattering, *etc*.) has been termed “inteingrated structural biology”. We present a combination of chemical shift perturbations (CSPs) with low-resolution cryoEM data, allowing the determination of structures of biomacromolecular complexes to high accuracy and precision.

Determining structures of complexes of (multidomain) proteins by X-ray crystallopgraphy has often proven to be challenging. Recently cryoEM has seen remarkable improvements in sensitivity, methodology and instrumentation, resulting in atomic structures, which are on par with X-ray structures in terms of resolution.^[1]^ Currently cryoEM is most readily applied for entities above a molecular weight (MW) of ca. 200-300 kDa.

NMR spectroscopy is the other major technique for obtaining structural data at atomic resolution. The development of transverse relaxation optimized spectroscopy (TROSY)^[2,3]^ in combination with uniform, high-level deuteration has pushed the MW limit of systems accessible to solution state NMR well above 100 kDa.^[4,5]^

In the interface region in which NMR reaches its upper and cryoEM its lower MW limit, synergies between the two techniques seem promising. In addition to the complementarity in terms of MW, the overall nature of the structural information delivered by both techniques can be used synergistically. While the accessible NMR observables at high MW are usually of local nature (commonly chemical shift perturbations, paramagnetic relaxation enhancements (PREs) and in favourable cases intermolecular NOEs), cryoEM is able to determine global information on the shape of the system under study.

The histidine-containing phosphoprotein (HPr) is a branching point intermediate in a phosphoryl-transfer signalling chain.^[6]^ It is able to transfer its phosphoryl group attached to its His15 residue to a number of sugar-specific acceptor proteins, so called IIA proteins. The individual structures of HPr and the glucose-specific IIA^Glc^ from *E.coli* were determined by X-ray crystallography at 2.0 and 1.5 Å resolution, respectively.^[7,8]^ The HPr-IIA^Glc^ complex could not be crystallized, probably due to its relatively low affinity.^[9]^ Measuring a set of 74 intermolecular NOEs between HPr and IIA^Glc^ and ca. 200 residual dipolar couplings (RDCs) of the complex allowed the determination of the complex structure by solution NMR.^[9]^ Rigid body docking of the individual crystal structures, followed by a restrained simulated annealing protocol giving torsional freedom to the interfacial side chain atoms in order to prevent clashes^[10,11]^ resulted in a well defined complex structure. The HPr-IIA^Glc^ complex has served as a test system for determining complex structures from very diverse sets of biophysical and biochemical parameters.

NOEs have high information content, but are difficult to obtain, particularly for larger systems. On the one hand this is due to the fact that complete chemical shift assignments are required to interpret NOESY spectra. On the other hand larger systems require high levels of deuteration in order to deliver NMR spectra of satisfactory quality, which is in direct conflict with measuring large numbers of intermolecular NOEs. It is therefore not surprising that alternatives to the use of NOEs have been sought for the determination of complex structures.

Several NMR observables have been successfully applied to the determination of biomacromolecular complex structures.^[12]^ The most readily accessible NMR observables are chemical shift perturbations (CSPs). Unfortunately, these do not contain any explicit distance information, but merely define the interaction surfaces of the components of a complex. They can be translated into a set of highly ambiguous interaction restraints (AIRs),^[13–15]^ which are, however, not usually sufficient to orient two interaction partners unambiguously relative to each other. Clore and Schwieters have shown that CSPs combined with a set of RDCs result in correct structures of the HPr-IIA^Glc^ complex^[16]^ (as verified by comparison with the structure determined from intermolecular NOEs and RDCs^[9]^). Our work described herein is based on this approach and we have used the published CSPs for the HPr-IIA^Glc^ complex in our structure calculations. We have substituted the use of RDCs for the application of EM maps of different resolution. We show in this communication that such data, even at low resolutions, together with CSPs is able to provide high quality structures of macromolecular complexes. The workflow of our proposed method is schematically depicted in Figure 1. We are aware that the molecular weight of the system under study is well below the currently accessible size for cryoEM. We have merely chose the system due to the fact that it is a particularly well studied system and with a wealth of published experimental NMR data.

**Figure 1.**
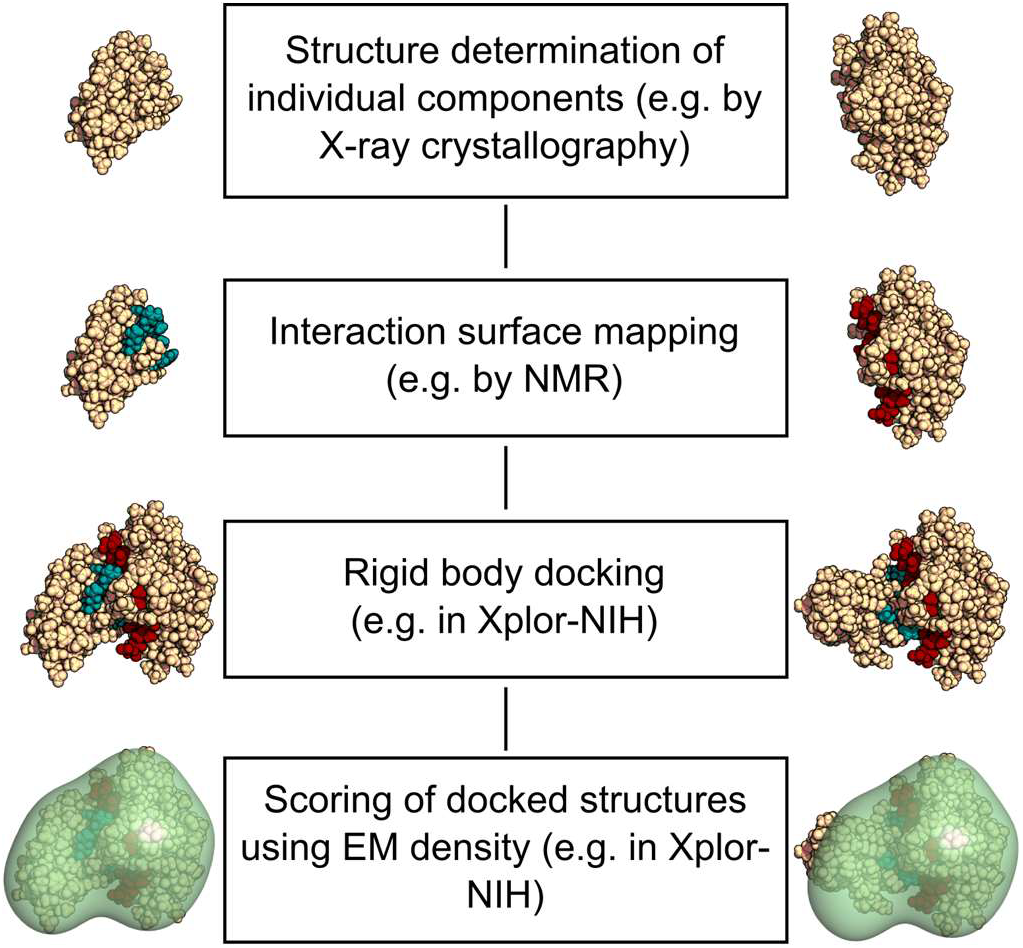
Structures of individual components of the complex are determined by traditional methods, *e.g.* X-ray crystallography. These are used as the starting point in our method. The interaction surface n the complex is mapped by chemical shift perturbation (CSP) mapping. CSP data are translated into highly ambiguous interaction restraints (AIRs). A large number of of complexes in agreement with these AIRs is calculated in Xplor-NIH. Complexes are then scored/sorted using low resolution cryoEM maps of the complex.

Titration of unlabelled HPr with ^15^N-labelled IIA^Glc^ and vice versa results in the mapping of a well defined interaction surface between the two subunits (Figure 2a). This information was translated into a set of highly ambiguous interaction restraints (AIRs) as previously described.^[16]^ The structure of the HPr-IIA^Glc^ complex as determined from intermolecular NOEs and residual dipolar couplings (RDCs)^[9]^ was used to generate artificial EM maps of different nominal resolutions (Figure 2b-d). The gradual loss of detail in going from a 10 Å map to lower resolutions is evident by eye.

**Figure 2.**
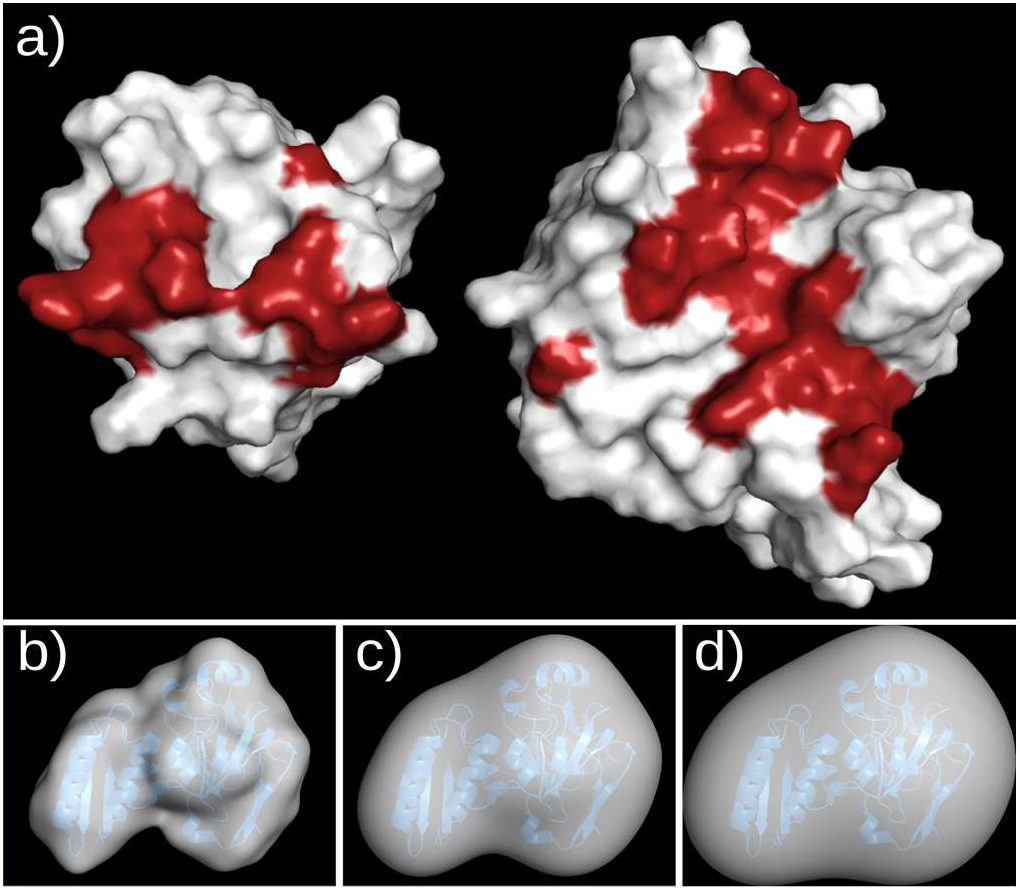
a) The contact surfaces of HPr (left) and IIA^Glc^ (right) as determined by chemical shift mapping are highlighted in red on a surface representation of the two proteins. For details on classification of the chemical shift perturbations see Clore and Schwieters, 2003.^[16]^ b-d) EM maps generated for the HPr-IIA^Glc^ complex (PDB code 1ggr) at different nominal resolutions. A ribbon representation in blue of the HPr-IIA^Glc^ complex is shown for clarity. Maps at 10 Å (b), 20 Å (c) and 30 Å (d) are shown.

The individual structures of HPr (2.0 Å resolution)^[7]^ and IIA^Glc^ (1.5 Å resolution)^[8]^ as determined by X-ray crystallography were used in a rigid body docking protocol described previously.^[16]^ The total energy (E_tot_) calculated in this protocol is a sum of Van der Waals (E_VdW_),^[17,18]^ side chain torsion angle potential (E_sc_),^[19]^ AIR (E_AIR_)^[13–15]^ and empirical radius of gyration (E_Rgyr_) terms.^[16,20]^ One hundred complex structures were generated, which showed a large structural diversity. Hierarchical clustering allows classifying the resulting structures based on structural similarity.^[21,22]^ We computed the pairwise RMSD between all 100 members of the ensemble and used this information to perform hierarchical clustering. Applying a 5 Å RMSD cutoff these structures fall into six distinct classes as visualized in Figure 3a.

**Figure 3.**
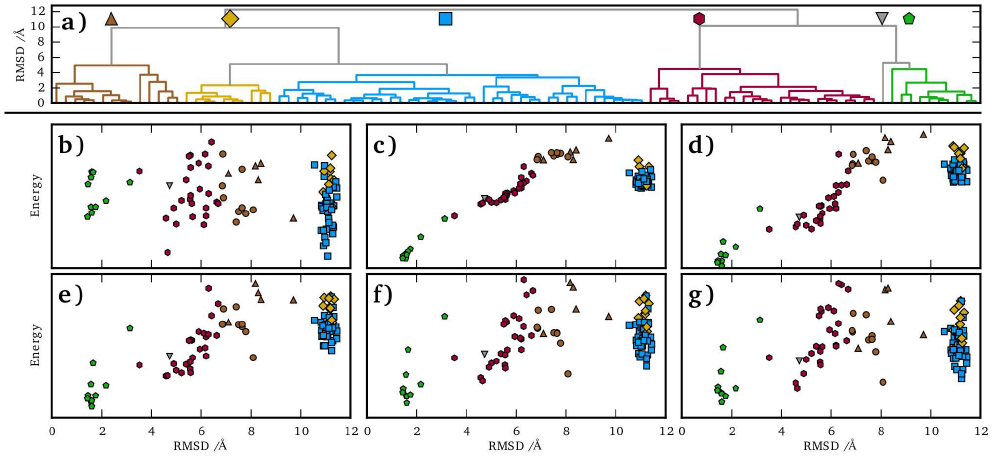
Structural clustering and correlations between the RMSD and the energy of structures docked using CSPs with and without inclusion of an EM map energy term. a) Clustering based on pairwise RMSD of complex structures generated with CSP data only using a 5 Å cutoff for the clustering. The different clusters are coloured arbitrarily. b-g) RMSDs of structures generated from docking vs. the 1ggr PDB structure. The symbols and colours correspond to the individual clusters in panel a) of the figure. b) Correlation of RMSD vs. the total energy of a given structure without the inclusion of EM data. c-g) Correlation of the RMSD with respect to the total energy including an EM map with resolution of 10 Å (c), 20 Å (d), 30 Å (e), 40 Å (f), and 50 Å (g).

CSPs alone are not sufficient for unambiguously orienting HPr relative to IIA^Glc^ (or vice versa) due to their low information content. This is reflected by a lack of correlation between E_tot_ of a complex structure calculated from CSPs alone versus the RMSD from the HPr-IIA^Glc^ complex structure deposited in the PDB (Figure 3b). Inspection of Figure 3b shows that only ca. 10% of the generated complexes have an RMSD <2 Å to the “true” structure of the complex and can thus be regarded as “correct” solutions. These “correct” structures are easily identified from the ensemble once EM densities of varying nominal resolution are used to score the structures obtained from docking HPr and IIA^Glc^. Surprisingly, EM densities at resolutions as low as 30-40 Å are sufficient to successfully differentiate correct from incorrect structures in the case of the HPr-IIA^Glc^ complex (Figure 3c-g). Only at resolutions above 40 Å EM maps start to fail at discriminating correct from incorrect structures.

Biomacromolecular interactions determine essentially every aspect of life. From a drug development perspective knowledge of the interaction surface of two molecules allows the possibility to screen for or rationally design potential inhibitors to a given pathway/network.^[23]^ Despite this importance the number of biomacromolecular complexes determined at high resolution to date remains quite low.^[24]^ Developing new methods for determining the structures of biomacromolecular complexes is therefore a vibrant field of research. We contribute to this field by showing that two molecules, whose individual crystal structures are known, can be docked onto one another by combining chemical shift perturbation data from NMR spectroscopy and densities from electron microscopy. EM densities of relatively modest resolution (up to 30-40 Å) are sufficient for the system under study. As such it can be envisaged that even negative staining EM, which results in much lower resolution maps as compared to cryoEM, will find useful application in our approach.

cryoEM has recently delivered a number of structures at atomic resolution (<4 Å). This was achieved on very large systems such as the ribosome, which are particularly well suited for cryoEM. Systems which are smaller, or in any other way less ideal for cryoEM (*e.g.* structurally heterogeneous, flexible, *etc*.), are expected to yield less detailed information. Even low resolution EM data proves successful in our docking approach. In combination with CSPs from NMR it can therefore still deliver high resolution complex structures in some cases. In the extreme case of a perfectly spherical complex, an EM map would not be able to deliver any additional information for scoring complexes derived from rigid body docking approaches. It is therefore conceivable that our approach is most promising for complexes showing a pronounced anisotropy. Similarly small-angle X-ray scattering (SAXS) has also been applied successfully for obtaining information on the global shape of complexes to help in the structure determination of biomacromolecular complexes.^[16,25,26]^ While SAXS is in general probably the more easily applicable method, cryoEM has the advantage that it can in principle deal with structural heterogeneous systems.^[27]^

Data derived from EM has previously been applied to structure calculations by both solution^[28]^ and solid state^[29,30]^ NMR. Including maps of resolutions ranging from 15 to 35 Å into structure refinement of RNAs has been shown to dramatically decrease the RMSD of the resulting structures.^[28]^ Also the popular docking suites HADDOCK^[15,31]^ and MODELLER^[32]^ have both recently also been extended to allow the incorporation of EM data.^[33,34]^ Our work has exclusively relied on CSPs for characterizing interface of the two partners. Utilizing perdeuteration and selective reprotonation of methyl groups, allows the measurement of CSPs for even very large complexes. The methodology herein is therefore applicable to many systems of biological interest. In principle any other method for mapping the interface in a complex should be suitable to replace CPSs in our approach. We envisage that hydrogen-deuterium exchange MS or chemical cross-linking detected by MS should be promising candidates for this. Methodology to incorporate such data into molecular docking has already been implemented in the above mentioned docking and modelling suites. Agreement of data obtained from different experimental techniques is of course a requirement that all integrated structural biology techniques rely on. As the extent of agreement may vary, it is important to verify it on a case by case basis.

Finally, as advances in NMR push to ever higher MW systems and concomitant improvements in cryoEM allow access to smaller systems, harnessing the synergies between these two techniques becomes of more and more interest. As CSPs are the most readily available NMR observable in larger systems and low resolution EM maps can be generated for even challenging systems, both experimental restraints can be considered to be relatively low hanging fruit. Thus we anticipate that the method presented herein will find a broad application in the near future.

## Experimental Section

The chemical shift perturbation (CSP) data upon complex formation of HPr and IIA^Glc^ used in this work has been published^[16]^ and are available as part of the Xplor-NIH software package.^[35]^ CSPs were converted to ambiguous interaction restraints (AIRs) and a conjoined rigid body/torsion angle dynamics protocol for structure calculation^[36,37]^ was used as described previously.^[16]^ The X-ray structures of HPr (PDB code 1poh)^[7]^ and IIA^Glc^ (PDB code 2f3g, molecule 2)^[8]^ were used in the rigid body docking protocol. EM densities for the HPr-IIA^Glc^ complex were generated from the structure deposited in the protein data bank (PDB code 1ggr)^[9]^ using the software packages Situs^[38]^ or Chimera.^[39]^ Methodology for handling EM densities in Xplor-NIH has recently been described^[28]^ and has been supplemented by in-house developed code (available from the authors upon request). All graphics were produced with PyMOL^[40]^ and plotting was carried out using the Python package matplotlib.^[41]^ All calculations were carried out on a desktop workstation running CentOS version 6.5 and Xplor-NIH version 2.39.^[35]^

Hierarchical clustering was performed with Scipy using the scipy.cluster.hierarchy class. A complete linking was conducted and a 5 Å RMSD cutoff was applied for clustering of the structures. The python code for performing the cluster analysis is available from the authors upon request.

## Acknowledgements

The authors thank Kevin J. Embrey (AstraZeneca) and Charles Schwieters (NIH) for helpful discussions and suggestions. This work was funded through the AstraZeneca Postdoc Programme.

## Entry for the Table of Contents

### COMMUNICATION

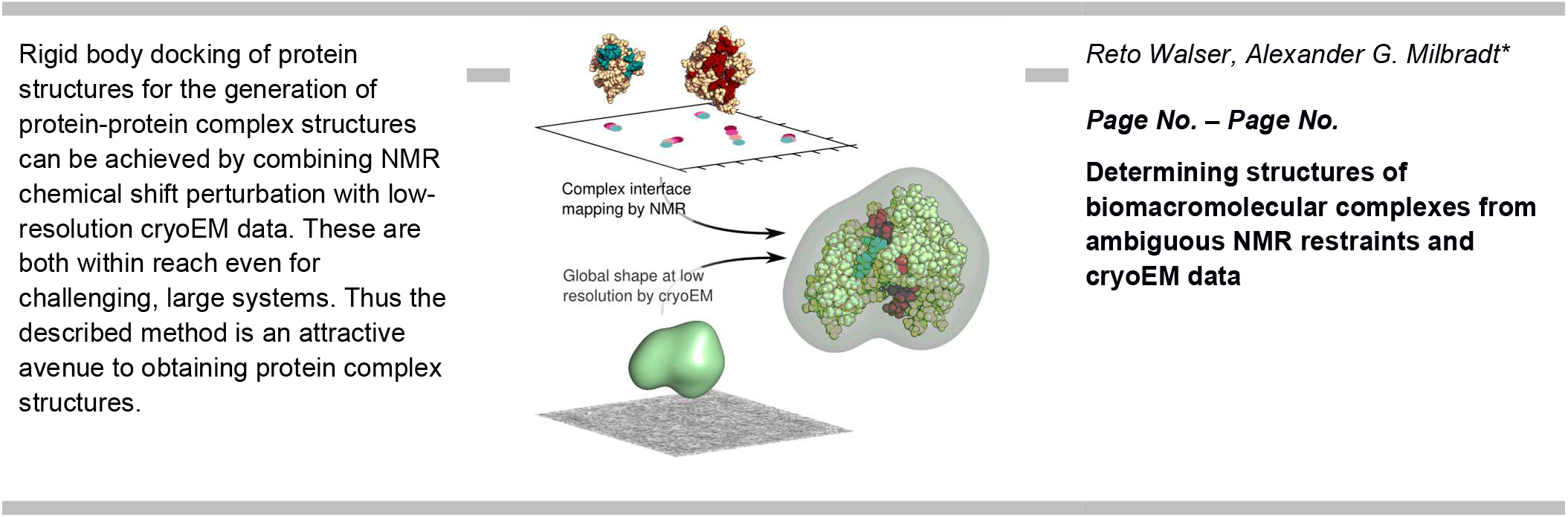

